# FAK-Dependent Focal Adhesion Remodeling Drives Arsenical Vesicant–Induced Skin Injury

**DOI:** 10.64898/2026.10.20.760501

**Authors:** Han Liu, Suman Suman, Alex Hoover, Ben Taubman, Shao-Yu Chen, Hening Lin, Rajesh Agarwal, Mohammad Athar, Xiaoyang Wu

## Abstract

Chemical vesicants cause severe skin injury characterized by inflammation, tissue destruction, and separation of the epidermis from the underlying dermis. However, the molecular mechanisms linking vesicant exposure to the disruption of epidermal adhesion remain poorly understood. Phenylarsine oxide (PAO), a commonly used lewisite analog, is a potent inducer of cutaneous blistering and a strong inhibitor of protein tyrosine phosphatases. Here, we show that PAO rapidly induces focal adhesion kinase (FAK) phosphorylation and downstream signaling in epidermal keratinocytes, accompanied by accelerated focal adhesion disassembly and molecular turnover. Genetic ablation of *FAK* stabilizes focal adhesions and markedly suppresses PAO-induced adhesion remodeling. Importantly, epidermal-specific deletion of *FAK* protects mouse skin from PAO-induced microvesication, inflammatory responses, and cell death. Pharmacological inhibition of FAK similarly preserves focal adhesion stability *in vitro* and attenuates PAO-induced skin injury *in vivo*, phenocopying the protective effects of genetic *FAK* deficiency. Together, our findings identify aberrant FAK activation as a critical signaling event linking arsenical vesicant exposure to focal adhesion failure and skin vesication, and suggest that targeting FAK may provide a therapeutic strategy to mitigate chemical vesicant-induced skin injury.

**Highlights:**

- First demonstration that focal adhesion kinase (FAK) is a critical molecular target of chemical vesicants whose dysregulation disrupts epidermal–dermal junctional integrity and initiates micro-vesicant injury.
- Establishes FAK-mediated junctional disruption as an early pathogenic event that triggers and amplifies subsequent inflammatory and programmed cell-death responses following vesicant exposure.
- Integrating *in vivo* genetic and pharmacological approaches, identifies FAK as a mechanistically validated and therapeutically actionable target for development of a mechanism-based medical countermeasure (MCM) against chemical vesicant injury.

## INTRODUCTION

Chemical warfare agents cause devastating acute and long-term health consequences^1,2^. Among these agents, chemical vesicants produce severe skin inflammation, tissue destruction, and blister formation that can lead to permanent disfigurement and disability. Cutaneous vesicants include mustards, arsenicals, and urticants^2–4^. Despite international restrictions on the production and use of chemical weapons, intentional deployment, accidental exposure, and remaining stockpiles continue to pose significant public health and security concerns^1,5,6^. However, effective mechanism-based medical countermeasures for vesicant-induced skin injury remain limited^3,7–9^. A better understanding of the molecular events^10^ that initiate tissue damage is therefore essential for developing therapeutic strategies that can prevent or reduce chemical vesication.

The skin epidermis forms the first line of defense against environmental injury^11,12^. Its barrier function depends on stable adhesion between neighboring keratinocytes and attachment of basal epidermal cells to the underlying basement membrane^13,14^. Cell-cell junctions maintain cohesion within the epidermis, whereas integrin-based focal adhesions connect keratinocytes to the extracellular matrix and coordinate the organization of the actin cytoskeleton^15,16^. These adhesion structures are highly dynamic and undergo continuous remodeling during cell migration, developmental morphogenesis, tissue repair, and tumor progression^15,17^. However, excessive or uncontrolled adhesion turnover can destabilize tissue architecture. Disruption of epidermal adhesion is a prominent feature of vesicant-induced injury and ultimately leads to dermal-epidermal separation and blister formation^9,18^. Nevertheless, the signaling mechanisms that connect chemical exposure to the rapid loss of epidermal adhesion remain poorly understood^10^.

Focal adhesion kinase (FAK) is a cytoplasmic tyrosine kinase that functions as a central regulator of integrin signaling and focal adhesion dynamics^19–23^. Following activation, FAK undergoes autophosphorylation at Tyr397 and recruits signaling proteins that regulate focal adhesion turnover, cytoskeletal remodeling, cell migration, survival, and inflammation. Important downstream components of this pathway include Src family kinases, paxillin, p130Cas, PI3K, MAPK, and NF-κB. FAK signaling has been extensively studied in tissue repair, fibrosis, inflammation, and cancer cell migration. However, whether aberrant FAK activation contributes to epithelial adhesion failure during chemical vesicant-induced skin injury remains unclear.

Phenylarsine oxide (PAO), a commonly used analog of the arsenical vesicant lewisite, is a potent protein tyrosine phosphatase inhibitor^24–26^. By disrupting the balance between protein phosphorylation and dephosphorylation, PAO may activate FAK signaling and alter focal adhesion organization^27,28^. We therefore hypothesized that PAO-induced FAK activation represents an early signaling event that accelerates focal adhesion turnover and promotes epidermal barrier disruption. To test this hypothesis, we employed a combination of live-cell imaging, biochemical analysis, mouse genetics, and pharmacological inhibition to examine the role of FAK in PAO-induced skin injury. Our results show that PAO rapidly activates FAK, leading to accelerated focal adhesion disassembly in epidermal keratinocytes. Genetic deletion of *FAK* stabilizes focal adhesions and protects mouse skin from PAO-induced dermal-epidermal separation, inflammation, and micro-vesicant (micro-blister) formation. Pharmacological inhibition of FAK similarly suppresses focal adhesion remodeling and attenuates PAO-induced skin injury *in vivo*. Together, our findings identify FAK-dependent adhesion remodeling as an important molecular mechanism underlying arsenical-induced skin vesication and suggest that targeting FAK may provide a therapeutic strategy for the development of medical countermeasures against chemical vesicants.

## RESULTS

### PAO promotes focal adhesion turnover in epidermal keratinocytes

Skin vesication is characterized by separation of the epidermis from the underlying dermal tissue, suggesting that disruption of cell-matrix adhesion is an important early event in this process^2–4^. Focal adhesions connect the actin cytoskeleton to the extracellular matrix and are critical for maintaining epidermal attachment^14,15^. To determine whether PAO directly affects cell/extracellular matrix adhesions, we first examined focal adhesion organization in wild-type primary mouse keratinocytes. Immunofluorescence microscopy showed prominent paxillin-positive focal adhesions, which were primarily localized at the cell periphery in untreated cells^29^. PAO treatment led to a marked reduction in focal adhesion size (Fig. 1A). Quantification of paxillin-positive structures confirmed that focal adhesions were substantially smaller in PAO-treated keratinocytes relative to untreated controls (Fig. 1B). Because FAK autophosphorylation at Tyr397 is a hallmark of FAK activation^20^, we next examined FAK phosphorylation by western blotting. PAO treatment increased FAK Tyr397 phosphorylation without an apparent change in total FAK abundance (Fig. 1C), providing biochemical evidence that PAO can activate FAK signaling in epidermal keratinocytes.

**Figure 1.**
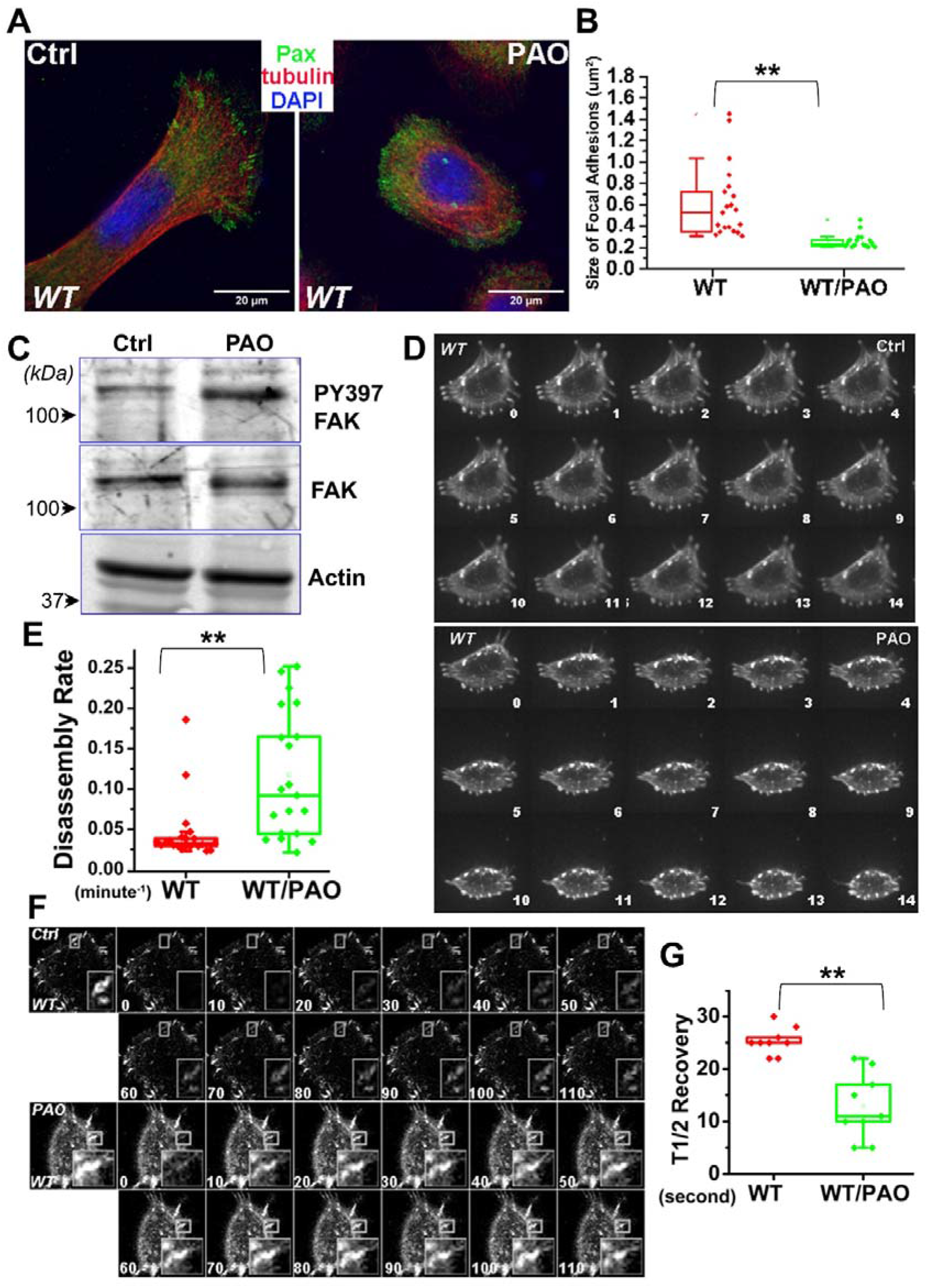
PAO activates FAK and accelerates focal adhesion remodeling in epidermal keratinocytes. (A) Primary wild-type (WT) keratinocytes treated with vehicle control (Ctrl) or PAO were stained for paxillin (Pax; green), α-tubulin (red), and DAPI (blue). Scale bars, 20 μm. (B) Quantification of the area of individual paxillin-positive focal adhesions. N=30. **: p<0.01 (Student’s t-test). (C) Western blot analysis of FAK Tyr397 phosphorylation (pY397 FAK), total FAK, and actin in control and PAO-treated keratinocytes. (D) Representative time-lapse montages of DsRed-Zyxin-labeled focal adhesions in control and PAO-treated WT keratinocytes; numbers indicate elapsed time in minutes. (E) Quantification of focal adhesion disassembly rates in control and PAO-treated WT keratinocytes. Box-and-whisker plots show individual measurements. N=50. **: p<0.01 (Student’s t-test). (F) Representative FRAP sequences of DsRed-Zyxin-labeled focal adhesions; numbers indicate seconds after photobleaching. (G) Quantification of focal adhesion disassembly rates and the half-time (T1/2) of fluorescence recovery after photobleaching (bottom) in control and PAO-treated WT keratinocytes. N=10. **: p<0.01 (Student’s t-test).

We further employed confocal videomicroscopy to trace and examine the behavior of individual focal adhesions. To visualize focal adhesions, we transfected keratinocytes with a plasmid encoding DsRed-Zyxin, a fluorescently labeled focal adhesion marker protein^29^. Representative examples of focal adhesion dynamics are shown as time-lapse montages in Figure 1D. During the interval of observation, focal adhesions in untreated keratinocytes underwent continual bouts of formation, maturation, and disassembly. By contrast, PAO treatment markedly accelerated focal adhesion turnover, with individual adhesions undergoing more rapid disassembly. Quantification of the kinetics of individual focal adhesions revealed a substantial increase in the disassembly rate following PAO treatment (Fig. 1E).

The changes in focal adhesion dynamics were further substantiated by fluorescence recovery after photobleaching (FRAP) experiments (Fig. 1F). Compared with untreated cells, PAO-treated keratinocytes exhibited a substantial decrease in the half-time of fluorescence recovery after bleaching (Fig. 1G), indicating increased molecular exchange within focal adhesions. Together, these data provide compelling evidence that PAO activates FAK and promotes the dynamic remodeling and turnover of focal adhesions in epidermal keratinocytes.

### FAK is required for PAO-induced focal adhesion remodeling

To determine the role of FAK in PAO-induced focal adhesion remodeling, we next examined cell/extracellular matrix adhesions after *FAK* deletion^30^. Immunofluorescence microscopy showed prominent paxillin-positive focal adhesions in FAK-null keratinocytes (Fig. 2A). In contrast to the marked reduction in focal adhesion size observed in PAO-treated wild-type cells, focal adhesions in FAK-null cells remained relatively large following PAO treatment. Quantification of paxillin-positive structures showed no substantial difference in focal adhesion size between untreated and PAO-treated FAK-null keratinocytes (Fig. 2B), suggesting that loss of *FAK* renders focal adhesions resistant to PAO-induced remodeling.

**Figure 2.**
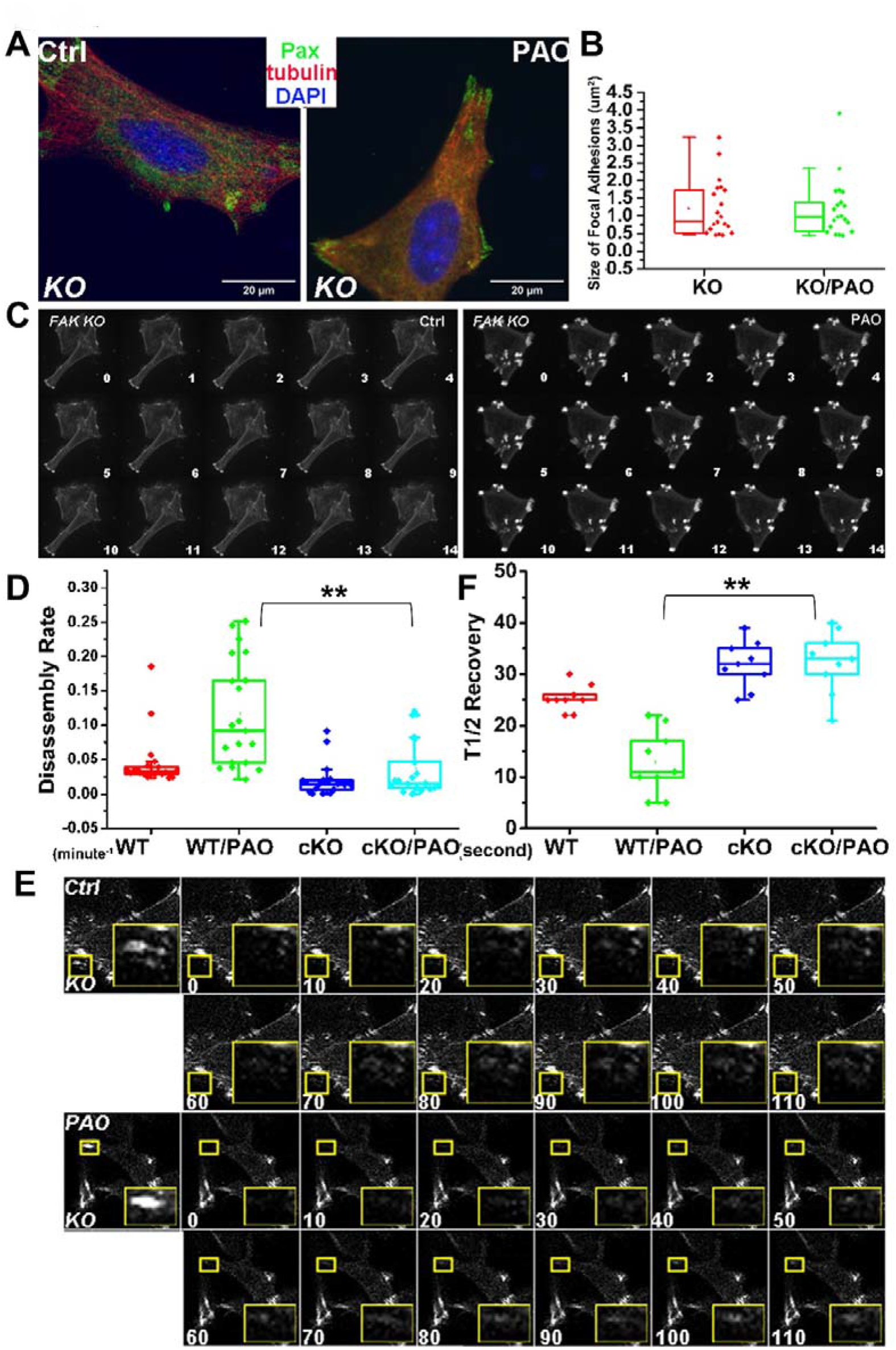
FAK is required for PAO-induced focal adhesion remodeling. (A) FAK-null keratinocytes treated with vehicle control (Ctrl) or PAO were stained for paxillin (Pax; green), α-tubulin (red), and DAPI (blue). Scale bars, 20 μm. (B) Quantification of the area of individual paxillin-positive focal adhesions. N=30. (C) Representative time-lapse montages of DsRed-Zyxin-labeled focal adhesions in control and PAO-treated *FAK*-null keratinocytes; numbers indicate elapsed time in minutes. (D) Quantification of focal adhesion disassembly rates in WT and FAK cKO keratinocytes with or without PAO. N=50. **: p<0.01 (Student’s t-test). (E) Representative FRAP sequences of DsRed-Zyxin-labeled focal adhesions in control and PAO-treated FAK-null keratinocytes; numbers indicate seconds after photobleaching. (F) Quantification of the half-time (T1/2) of fluorescence recovery. Box-and-whisker plots show individual measurements. N=10. **: p<0.01 (Student’s t-test)

We further employed confocal videomicroscopy to trace and examine the behavior of individual focal adhesions in *FAK*-null keratinocytes expressing DsRed-Zyxin. Representative examples of focal adhesion dynamics are shown as time-lapse montages in Figure 2C. During the interval of observation, focal adhesions in *FAK*-null cells were largely static and exhibited minimal turnover. Importantly, PAO treatment failed to promote rapid focal adhesion disassembly in the absence of FAK. Quantification of the kinetics of individual focal adhesions revealed that disassembly rates remained low in *FAK*-null cells following PAO treatment (Fig. 2D).

The defects in focal adhesion dynamics were further substantiated by FRAP experiments (Fig. 2E). Compared with wild-type cells, *FAK*-null keratinocytes displayed a substantial increase in the half-time of fluorescence recovery after bleaching, consistent with reduced molecular exchange within focal adhesions. Whereas PAO shortened the recovery half-time in wild-type keratinocytes, it had little effect on fluorescence recovery in *FAK*-null cells (Fig. 2F). Together, these data provide compelling evidence that FAK is required for PAO-induced focal adhesion remodeling and that loss of *FAK* stabilizes cell/extracellular matrix adhesions in epidermal keratinocytes.

### FAK mediates PAO-induced disruption of cell-cell junctions

To determine whether PAO affects tight-junction dynamics, we performed FRAP analysis of EGFP-ZO-1 at cell/cell contacts. Representative time-lapse images of PAO-treated wild-type and *FAK*-null keratinocytes are shown in Figure 3A, with the corresponding untreated controls shown in Supplementary Figure 1. PAO treatment accelerated fluorescence recovery in wild-type keratinocytes, as indicated by a substantially reduced recovery half-time (Fig. 3B). In contrast, *FAK*-null cells exhibited slower recovery kinetics and were relatively insensitive to PAO. These findings indicate that PAO increases ZO-1 exchange at cell/cell contacts and promotes tight-junction disassembly in a FAK-dependent manner.

**Figure 3.**
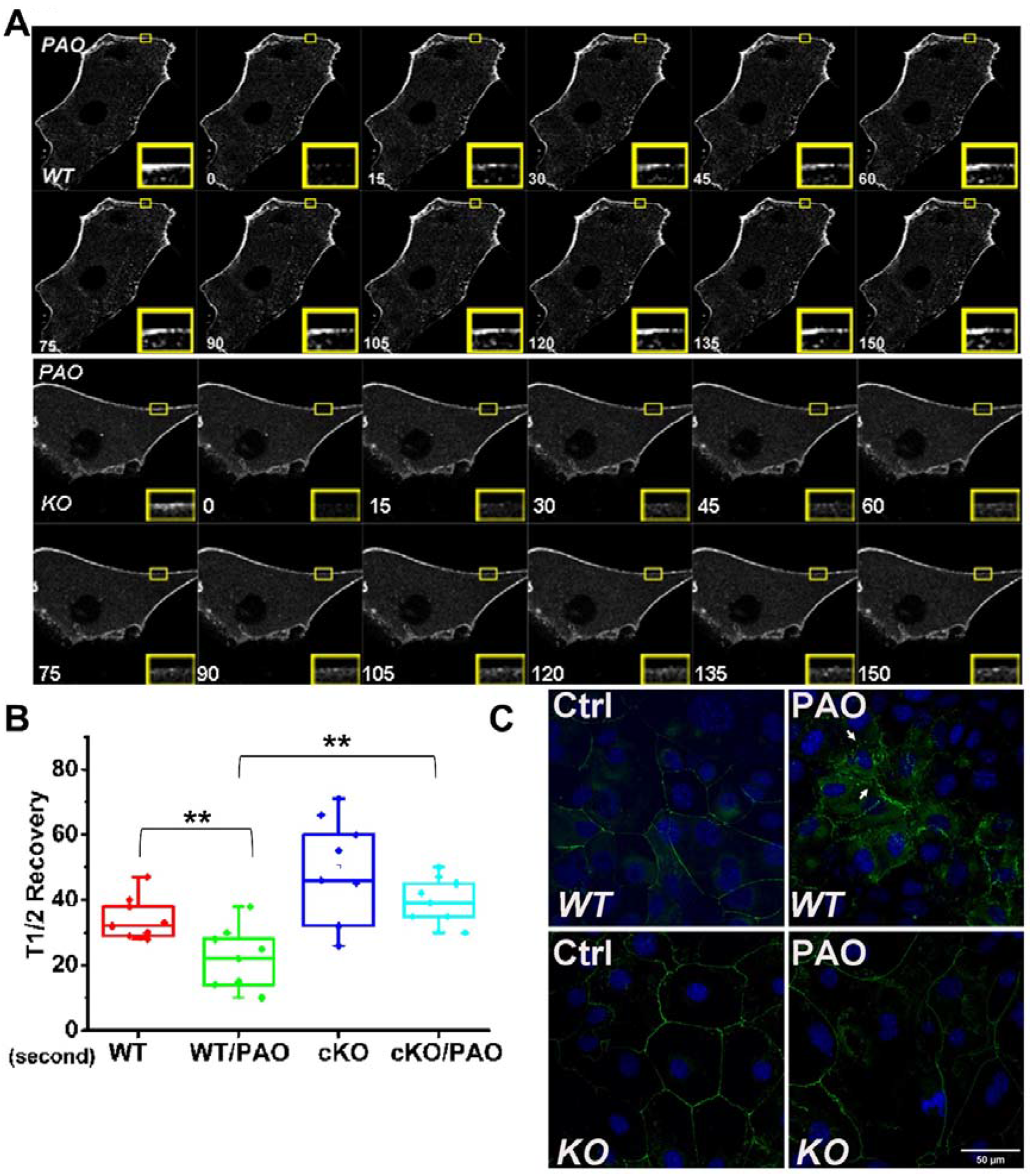
Loss of FAK stabilizes cell-cell junctions following PAO treatment. (A) Representative FRAP sequences of EGFP-ZO1 at cell-cell junctions in PAO-treated WT and FAK-null keratinocytes; numbers indicate seconds after photobleaching. Enlarged regions are indicated by yellow boxes. (B) Quantification of the half-time (T1/2) of fluorescence recovery in WT and FAK cKO keratinocytes with or without PAO. Box-and-whisker plots show individual measurements. N=10. **: p<0.01 (Student’s t-test). (C) Immunofluorescence staining of E-cadherin (green) and DAPI (blue) in WT and FAK-null keratinocytes treated with vehicle control (Ctrl) or PAO. Arrows indicate intracellular ZO-1 accumulation following PAO treatment. Scale bar, 50 μm.

We next examined ZO-1 localization by immunofluorescence microscopy. In untreated wild-type cells, ZO-1 was predominantly localized at cell/cell borders, forming continuous junctional structures. Following PAO treatment, junctional ZO-1 became disorganized, with a marked accumulation of ZO-1 within the cytoplasm. By contrast, FAK-null keratinocytes maintained relatively continuous ZO-1-positive junctions and showed less intracellular ZO-1 accumulation following PAO treatment (Fig. 3C). Together, these results demonstrate that PAO promotes FAK-dependent tight-junction disassembly and redistribution of ZO-1 from cell/cell contacts to intracellular compartments.

### Epidermal loss of FAK protects skin from PAO-induced vesication

To determine the role of FAK in chemical-induced skin injury *in vivo*, we generated epidermal-specific *FAK* conditional knockout (cKO) mice by crossing *FAK*^flox/flox^ mice^30^ with *K14-Cre* transgenic mice^31^. FAK cKO animals are viable and exhibit relatively normal skin architecture (Fig. 4A), allowing us to assess their response to PAO exposure. A single dose of PAO was topically applied to the dorsal skin of wild-type and *FAK* cKO mice^9^. PAO treatment induced a rapid and prominent erythema response in wild-type skin (Fig. 4B). By contrast, FAK cKO skin exhibited a markedly reduced gross response to PAO (Fig. 4A). Quantification using erythema scores confirmed that the cutaneous response to PAO was substantially attenuated in *FAK* cKO mice at eight hours after treatment (Fig. 4C).

**Figure 4.**
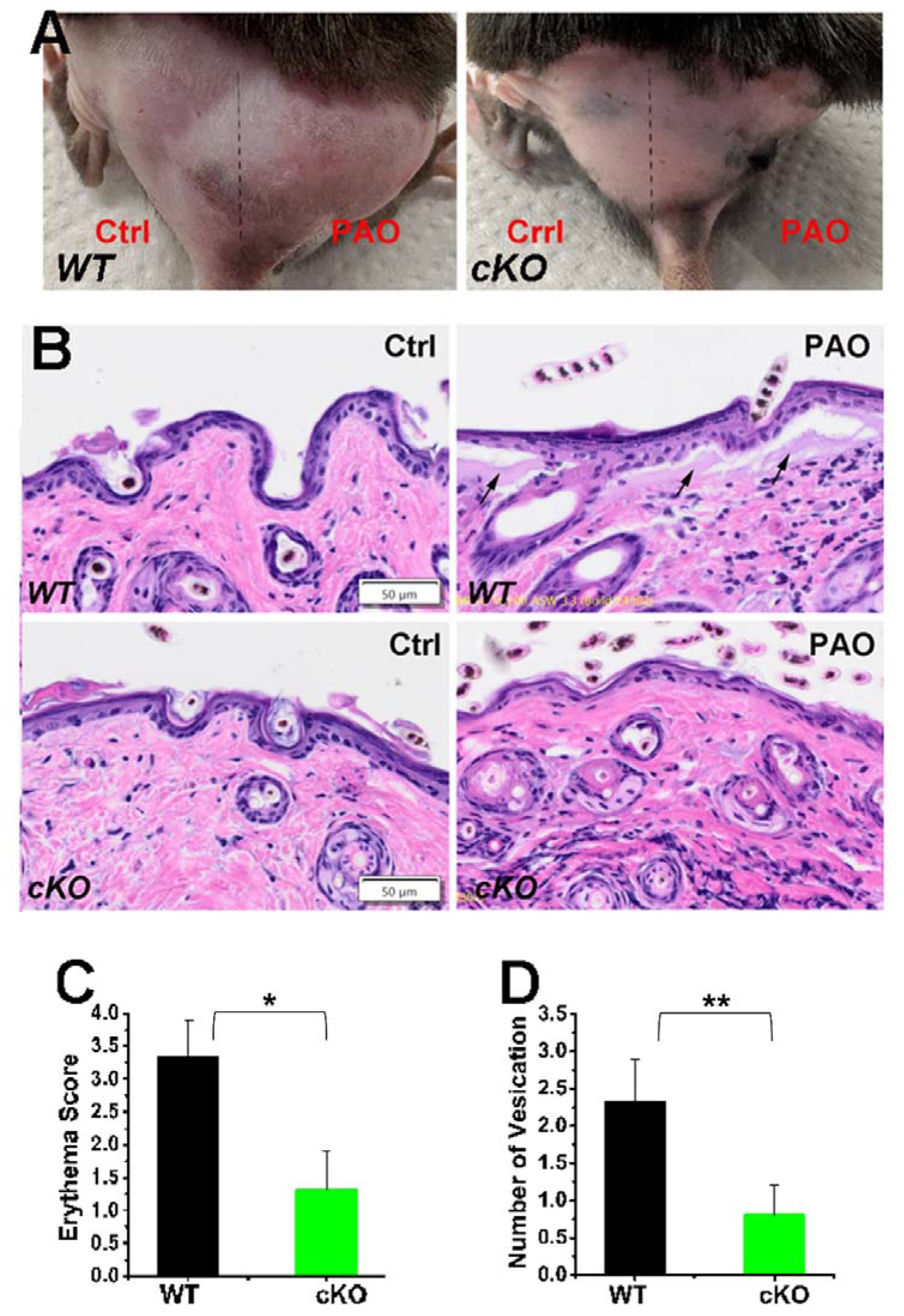
Epidermal deletion of *FAK* protects skin from PAO-induced vesication. (A) Representative gross appearance of dorsal skin from WT and epidermal-specific *FAK* conditional knockout (cKO) mice following vehicle control (Ctrl) or PAO treatment. The dashed line marks the boundary between treatment areas. (B) Representative hematoxylin and eosin-stained skin sections. Arrows indicate dermal-epidermal separation and microvesication formation in PAO-treated WT skin. Scale bars, 50 μm. (C) Quantification of erythema scores 8 hours after PAO exposure in WT and cKO mice. Error bars represent mean ± SD; n = 5 mice per group. **: p<0.01 (Student’s t-test). (D) Quantification of vesication events 8 hours after PAO exposure in WT and cKO mice. Error bars represent mean ± SD; n = 50 (fields). *: p<0.05, **: p<0.01 (Student’s t-test).

We next also examined skin histology for vesication following PAO exposure. Untreated wild-type and *FAK* cKO skin displayed intact epidermal architecture and attachment to the underlying dermis. PAO treatment caused extensive tissue disruption in wild-type skin, characterized by separation of the epidermis from the dermis and the formation of multiple microvesications. In contrast, epidermal–dermal attachment was largely preserved in PAO-treated *FAK* cKO skin, with fewer and less extensive areas of tissue separation (Fig. 4B). Quantification of individual vesication events confirmed a substantial reduction in microvesications formation following epidermal deletion of FAK (Fig. 4D).

Together, these data provide compelling *in vivo* evidence that FAK contributes to PAO-induced disruption of epidermal–dermal attachment and that loss of epidermal FAK protects skin from chemical vesication.

### FAK deficiency attenuates PAO-induced inflammatory signaling and cell death

To further characterize the tissue response to PAO, we first examined the expression of inflammatory markers in wild-type and *FAK* cKO skin. Immunohistochemical staining revealed a marked increase in COX-2 expression following PAO treatment in wild-type skin. By contrast, PAO-induced COX-2 expression was substantially reduced in *FAK* cKO skin (Fig. 5A). Quantification of the COX-2-positive area confirmed attenuation of this inflammatory response following epidermal deletion of FAK (Supplementary Fig. 2A).

**Figure 5.**
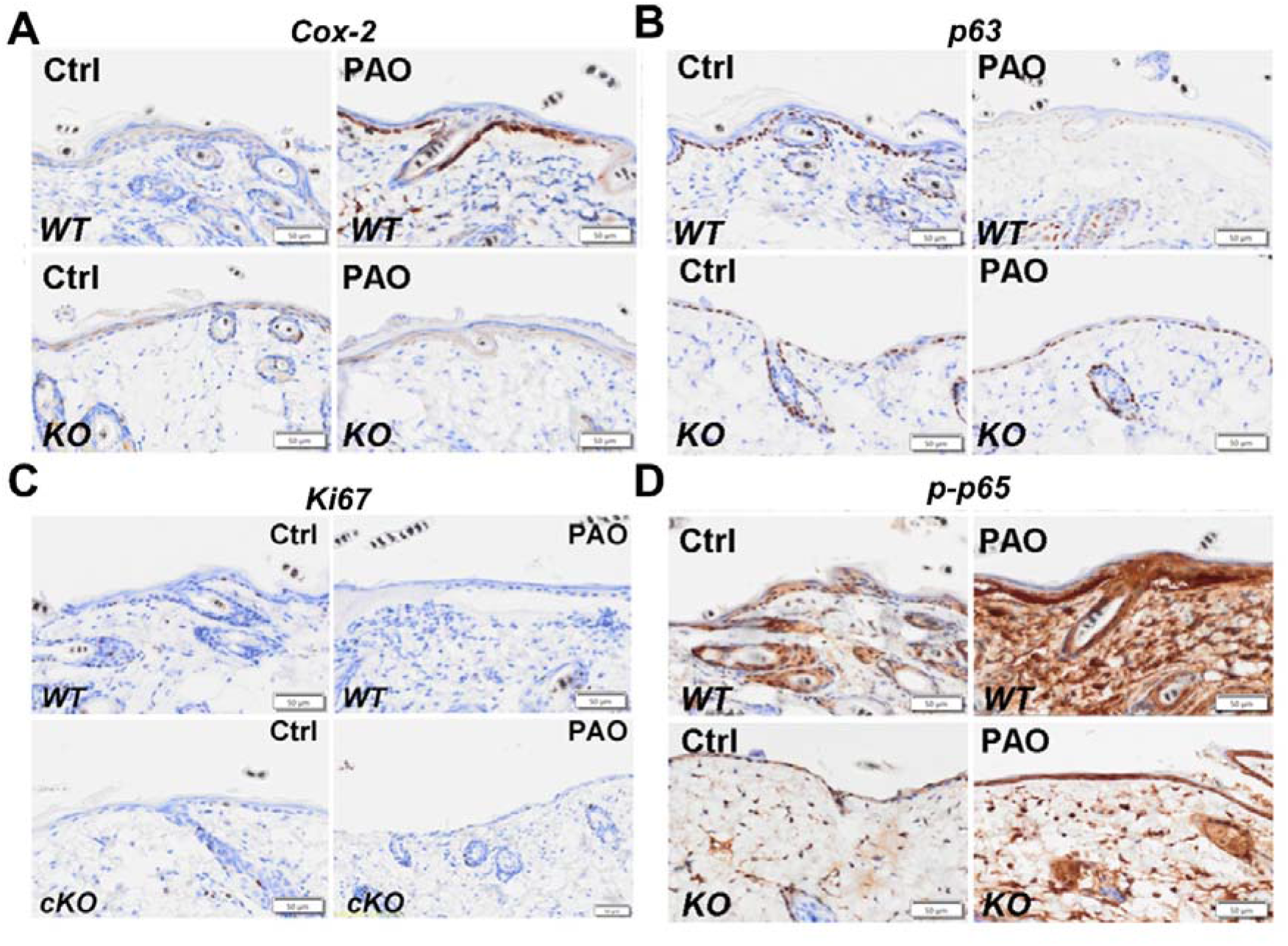
Epidermal FAK deficiency attenuates PAO-induced inflammatory signaling and preserves basal keratinocytes. Representative immunohistochemical staining of skin from WT and FAK cKO mice following vehicle control (Ctrl) or PAO treatment for (A) COX-2, (B) p63, (C) Ki67, and (D) phosphorylated NF-κB p65 (p-p65). Scale bars, 50 μm. Corresponding quantitative analyses are shown in Supplementary Figure 2.

Additional immunohistochemical analyses were performed to characterize the epidermal response to PAO. p63 is a master regulator controlling epidermal morphogenesis and stemness^57,58^. In untreated skin, p63-positive keratinocytes were primarily present within the basal epidermal layer. PAO treatment markedly reduced the number of p63-positive cells in wild-type skin, whereas p63 staining was largely preserved in *FAK* cKO skin following PAO exposure (Fig. 5B and Supplementary Fig. 2B). Ki67 staining remained relatively low across the examined conditions, suggesting that differences in the tissue response were not associated with a major increase in keratinocyte proliferation at this time point (Fig. 5C and Supplementary Fig. 2C).

We next examined activation of NF-κB signaling by immunostaining for phosphorylated NF-κB p65. PAO treatment induced a prominent increase in phospho-p65 staining in wild-type skin. Although phospho-p65-positive nuclei were also increased in PAO-treated *FAK* cKO skin, the response was reduced relative to PAO-treated wild-type skin (Fig. 5D and Supplementary Fig. 2D). Together, these findings suggest that loss of epidermal *FAK* attenuates PAO-induced activation of the NF-κB-COX-2 inflammatory pathway.

Finally, we examined PAO-induced cell death by TUNEL staining. PAO treatment produced a robust increase in TUNEL-positive cells in wild-type skin. FAK cKO skin exhibited lower TUNEL positivity following PAO treatment, although untreated FAK cKO skin showed a higher basal level of TUNEL staining than untreated wild-type skin (Supplementary Fig. 3). Thus, the relative increase in cell death following PAO exposure was substantially reduced by epidermal deletion of FAK. Together, these data indicate that the protection from vesication observed in *FAK* cKO skin is accompanied by attenuation of inflammatory signaling, preservation of the basal keratinocyte compartment, and reduced PAO-induced cell death.

### Pharmacological inhibition of FAK suppresses PAO-induced focal adhesion remodeling

To determine whether pharmacological inhibition of FAK can recapitulate the effects of genetic *FAK* deletion, wild-type keratinocytes were treated simultaneously with PAO and VS-6063 (defactinib), a FAK/PYK2 inhibitor ^32,33^. Immunofluorescence microscopy showed that keratinocytes treated with VS-6063 and PAO maintained large and prominent paxillin-positive focal adhesions (Fig. 6A). In contrast to the marked reduction in focal adhesion size induced by PAO alone, focal adhesions remained relatively large in the presence of the FAK inhibitor. Quantification of paxillin-positive structures confirmed that VS-6063 prevented the PAO-induced decrease in focal adhesion size (Fig. 6B).

**Figure 6.**
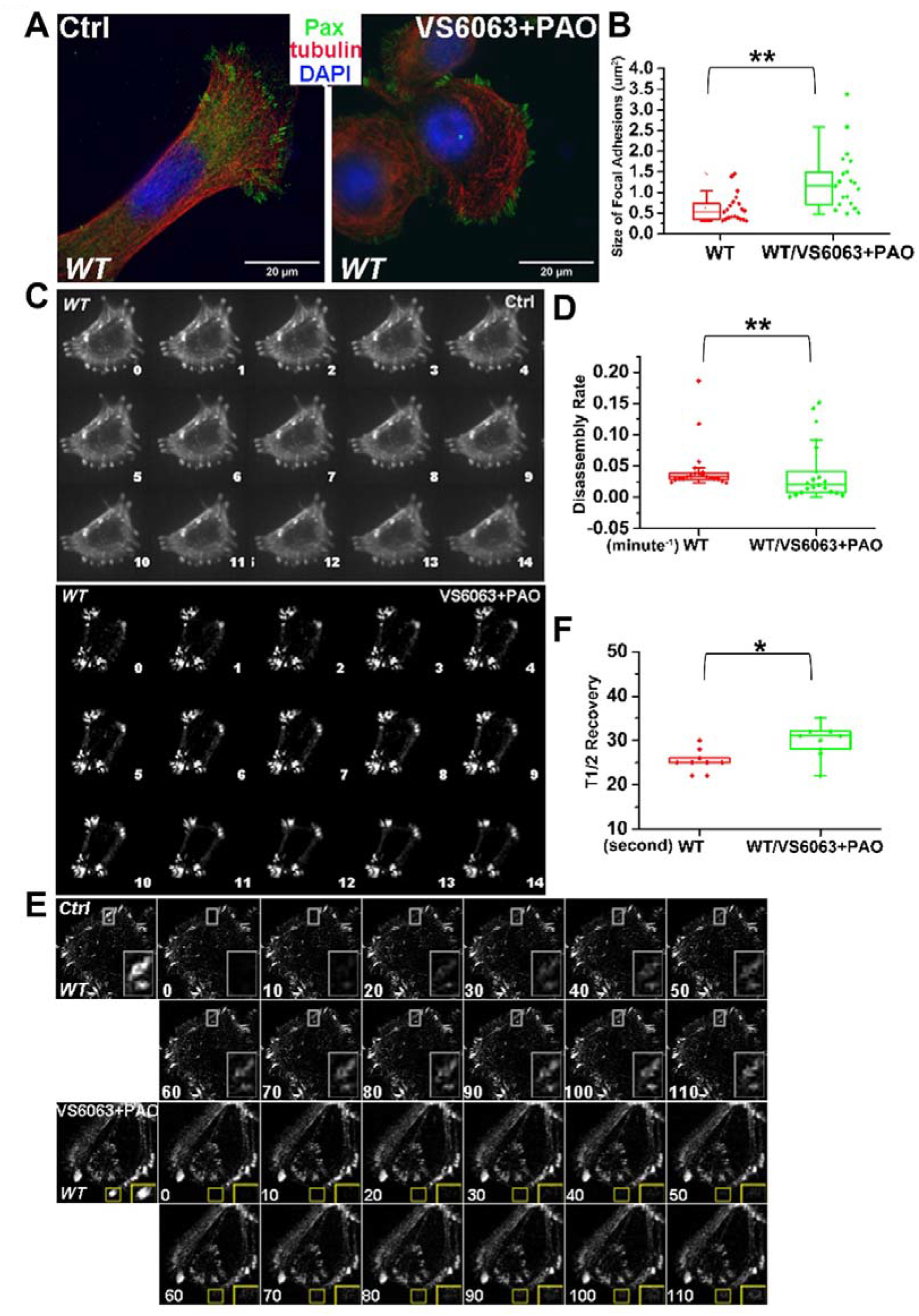
Pharmacological inhibition of FAK suppresses PAO-induced adhesion remodeling. WT keratinocytes were pretreated with VS-6063 before PAO exposure. (A) Cells were stained for paxillin (Pax; green), α-tubulin (red), and DAPI (blue). Scale bars, 20 μm. (B) Quantification of the area of individual paxillin-positive focal adhesions. N=30. **: p<0.01 (Student’s t-test). (C) Representative time-lapse montages of DsRed-Zyxin-labeled focal adhesions in control and VS-6063 plus PAO-treated cells; numbers indicate elapsed time in minutes. (D) Quantification of focal adhesion disassembly rates. N=50. **: p<0.01 (Student’s t-test). (E) Representative FRAP sequences of DsRed-Zyxin-labeled focal adhesions; numbers indicate seconds after photobleaching. (F) Quantification of the half-time (T1/2) of fluorescence recovery. N=10. *: p<0.05 (Student’s t-test).

We further employed confocal videomicroscopy to trace the behavior of individual focal adhesions in living keratinocytes expressing DsRed-Zyxin. Representative time-lapse montages are shown in Figure 6C. Focal adhesions in cells treated with VS-6063 and PAO remained relatively stable throughout the interval of observation. Unlike PAO treatment alone, which markedly accelerated focal adhesion turnover, FAK inhibition prevented the rapid dissolution of individual adhesions. Quantification showed that focal adhesion disassembly rates remained low following combined VS-6063 and PAO treatment (Fig. 6D).

The stabilization of focal adhesions by VS-6063 was further substantiated by FRAP analysis (Fig. 6E). Whereas PAO treatment shortened the half-time of fluorescence recovery, cells treated with VS-6063 before PAO exposure exhibited prolonged recovery kinetics. Quantification showed that the recovery half-time was restored toward that observed in untreated wild-type cells (Fig. 6F), indicating that FAK inhibition suppresses the PAO-induced increase in molecular exchange within focal adhesions.

Together, these data provide compelling evidence that pharmacological inhibition of FAK stabilizes focal adhesions and suppresses PAO-induced focal adhesion remodeling. VS-6063 also preserved junctional ZO-1 localization in PAO-treated wild-type keratinocytes (Supplementary Fig. 4), indicating that FAK inhibition protects both cell/extracellular matrix adhesions and cell/cell junctions. The similarity between the pharmacological and genetic results further supports an essential role for FAK in mediating the effects of PAO on epidermal adhesion. These studies also confirm the earlier observations showing impact of arsenicals in disrupting tight and adherents junctions forming proteins (ZO1 & ZO2).

### Pharmacological inhibition of FAK protects against PAO-induced skin vesication *in vivo*

To determine whether pharmacological inhibition of FAK protects against PAO-induced skin injury *in vivo*, wild-type mice were administered VS-6063 30 minutes after topical PAO exposure. PAO induced a prominent erythema response in vehicle-treated skin. By contrast, mice treated with VS-6063 exhibited a markedly reduced gross response to PAO (Fig. 7A). Quantification of erythema scores confirmed that post-exposure pharmacological inhibition of FAK attenuated the cutaneous response to PAO (Fig. 7C).

**Figure 7.**
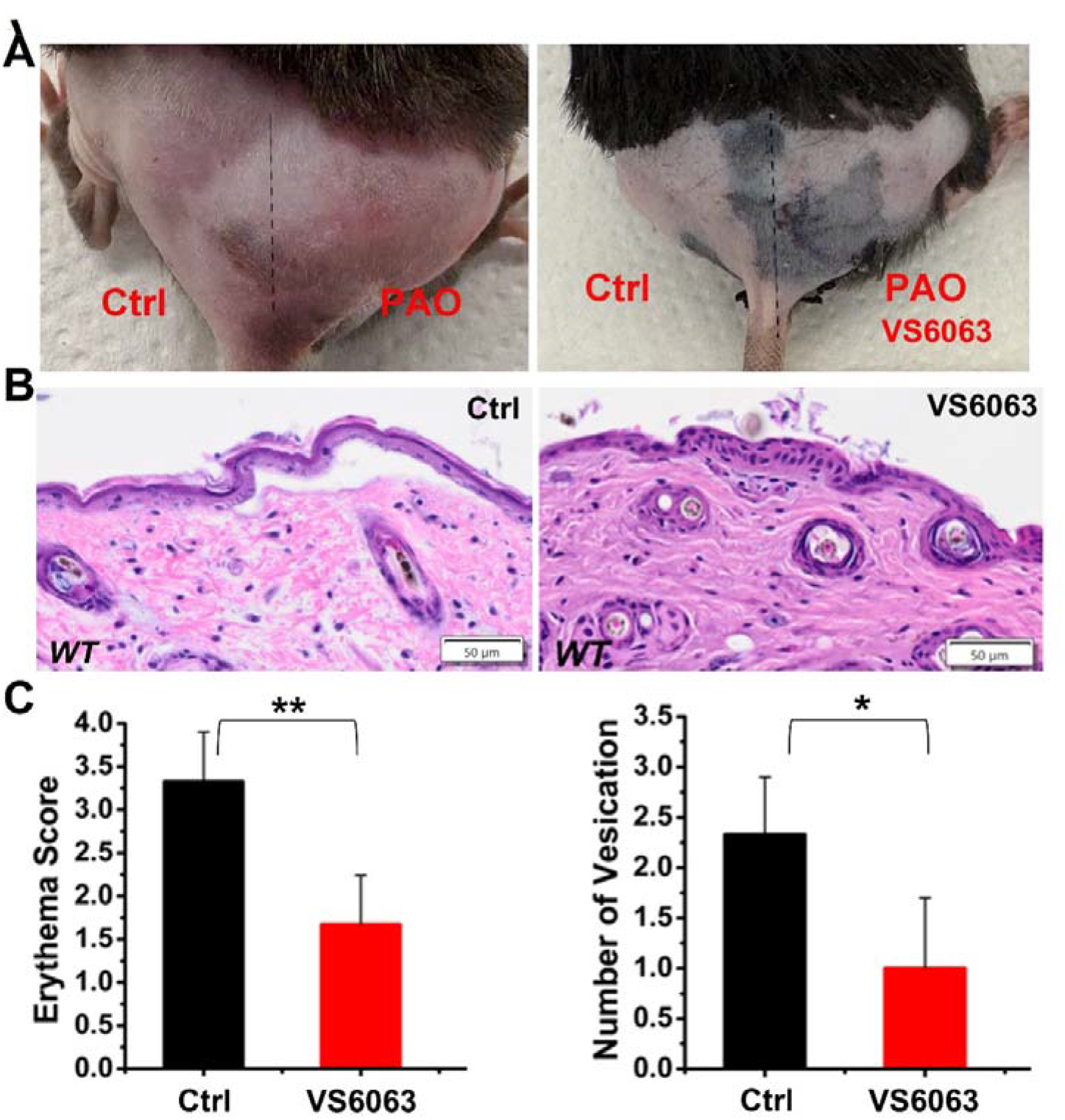
Post-exposure FAK inhibition protects against PAO-induced skin vesication. WT mice received VS-6063 30 minutes after topical PAO exposure. (A) Representative gross appearance of vehicle- and VS-6063-treated dorsal skin. The dashed line marks the boundary between treatment areas. (B) Representative hematoxylin and eosin-stained sections of PAO-exposed skin treated with vehicle control (Ctrl) or VS-6063. Scale bars, 50 μm. (C) Quantification of erythema scores and vesication events. Error bars represent mean ± SD; n = 5 mice per group for erythema score, n= 50 (fields) for quantification of vesication. *: p<0.05, **: p<0.01 (Student’s t-test).

Histological analysis further demonstrated the protective effect of VS-6063. PAO-treated skin exhibited disruption of normal tissue architecture, separation of the epidermis from the underlying dermis, and formation of multiple microvesications. In contrast, PAO-exposed skin treated with VS-6063 showed better preservation of epidermal-dermal attachment and fewer areas of tissue separation (Fig. 7B). Quantification confirmed a substantial reduction in the number of vesication events following FAK inhibitor treatment (Fig. 7C).

Together, these data demonstrate that post-exposure pharmacological inhibition of FAK protects against PAO-induced skin vesication in vivo. The ability of VS-6063 to phenocopy the protective effects of epidermal FAK deletion provides additional evidence that aberrant FAK activity contributes to chemical-induced skin injury and supports further evaluation of FAK inhibitors as potential medical countermeasures against arsenical vesicants.

## DISCUSSION

Chemical vesicants cause rapid disruption of epidermal integrity, but the early signaling events connecting chemical exposure to skin blistering remain incompletely understood^2–4^. In the present study, we identify FAK-dependent adhesion remodeling as an important mechanism underlying PAO-induced skin injury. PAO increased FAK Tyr397 phosphorylation, reduced focal adhesion size, accelerated focal adhesion disassembly, and increased the molecular exchange of focal adhesion proteins in epidermal keratinocytes. PAO also disrupted cell/cell junctions, suggesting a broader loss of epithelial adhesion. Genetic deletion of *FAK* stabilized these adhesion structures and protected mouse skin from PAO-induced erythema, epidermal-dermal separation, inflammation, and cell death. Importantly, pharmacological inhibition of FAK with VS-6063 recapitulated the protective effects of FAK deficiency in vitro and protected skin when administered after PAO exposure in vivo. Together, our findings establish aberrant FAK activation and FAK-dependent adhesion remodeling as a mechanistic link between arsenical exposure and epidermal barrier failure.

A central question arising from this study is how PAO increases FAK activity. FAK phosphorylation and focal adhesion turnover are controlled by a dynamic balance between protein tyrosine kinases and phosphatases^19–23^. PAO is a thiol-reactive trivalent arsenical and a potent inhibitor of protein tyrosine phosphatases^27,28^. The observed increase in FAK Tyr397 phosphorylation is consistent with a model in which PAO modifies critical cysteine residues within one or more phosphatases that normally restrain FAK signaling, thereby shifting the balance toward FAK activation. Several phosphatases, including PTP1B, SHP2, and PTP-PEST, have been implicated in the regulation of FAK, paxillin, and p130Cas^19–23^. PTP-PEST (PTPN12) is a well-established FAK Tyr397 phosphatases. Using AlphaFold, we predicted the possible binding interaction between human FAK1 and PTP-PEST. Interestingly, the prediction put the pTyr397 of FAK1 right into the active site of PTP-PEST, next to the catalytic Cys231 residue (Supplementary Figure 5). Furthermore, another Cys residue, Cys235, is adjacent to Cys231. PAO is known to target vicinal cysteines. Thus, it is likely that Cys231 and Cys235 of PTP-PEST are covalently targeted by PAO, leading to the inhibition of PTP-PEST and increase in FAK pTyr397. PAO-induced oxidative stress may further enhance this response because the catalytic cysteine residues of protein tyrosine phosphatases are sensitive to oxidation. Future studies using phosphatase-activity profiling and phosphatase-specific genetic approaches will be essential for identifying the direct molecular target responsible for FAK activation.

PAO can modify many proteins containing appropriately positioned thiol groups ^34^. Thus, the requirement for FAK does not necessarily indicate that FAK itself is the direct target of PAO. PAO may affect upstream phosphatases, integrin-associated proteins, small GTPases, or other components of the cytoskeletal machinery. These changes could cooperate to alter mechanical tension and activate FAK at adhesion sites. Time-resolved analysis of FAK phosphorylation, phosphatase activity, and focal adhesion turnover will help define the sequence of events connecting PAO exposure to adhesion failure.

Whether these findings can be extended to other arsenical vesicants is an important question. PAO is commonly used as a surrogate for lewisite, and both agents produce similar cutaneous pathogenetic responses, including erythema, inflammation, epidermal cell death, and epidermal–dermal separation. Because PAO and lewisite are both trivalent arsenicals that react with protein thiols, the FAK-dependent mechanism identified here may also contribute to lewisite-induced vesication. Direct validation using lewisite and other structurally distinct warfare arsenicals will be required to determine whether FAK represents a shared therapeutic target for arsenical skin injury.

Our study also highlights the coordinated regulation of cell/ECM adhesions and cell/cell junctions during vesicant-induced epithelial injury. These adhesion structures are physically and functionally connected through the cytoskeletal network^14,16^. PAO accelerated the dynamics of both systems, whereas loss of FAK stabilized them. We therefore posit that aberrant FAK activity induces a broader mechanical destabilization of the epithelial sheet rather than simply causing the dissolution of individual focal adhesions. Failure of cell/ECM attachment may redistribute cytoskeletal tension to cell/cell junctions, further weakening epidermal integrity and promoting tissue separation.

In addition to preserving epidermal adhesion, loss of *FAK* attenuated PAO-induced NF-κB activation and subsequent enhancement of COX-2 and cell death genes expression. FAK may directly promote inflammatory signaling following PAO exposure. Alternatively, the reduced inflammatory response may occur secondarily because preservation of epidermal architecture limits cell detachment, cell death, and the release of damage-associated molecules. These mechanisms are not mutually exclusive. The preservation of p63-positive basal keratinocytes, together with the relatively low Ki67 response, suggests that protection is not caused by compensatory proliferation. Instead, *FAK* deficiency appears to preserve the existing epidermal compartment and prevent the progression from adhesion failure to cell death and inflammation.

The protective effects of VS-6063 have important translational implications^32,33^. Existing FAK inhibitors may provide a direct path toward developing medical countermeasures for arsenical-induced skin injury. Importantly, administration of VS-6063 after PAO exposure reduced erythema and vesication, supporting the feasibility of post-exposure intervention. However, FAK also contributes to normal keratinocyte migration and wound repair. The timing, route, and duration of treatment will therefore be critical. Local administration may suppress the early wave of pathological FAK activity while limiting interference with subsequent tissue repair. An important next step will be to define the therapeutic window and determine how long after arsenical exposure FAK inhibition remains protective.

In closing, our findings provide critical insights into the molecular machinery underlying arsenical-induced skin vesication. We propose that PAO perturbs thiol-sensitive signaling to promote aberrant FAK activity, excessive remodeling of cell/ECM and cell/cell adhesions, and subsequent epidermal barrier failure. Genetic deletion or pharmacological inhibition of FAK interrupts this process and preserves skin integrity. These findings provide a rationale for further evaluating FAK as a therapeutic target for arsenical-induced cutaneous injury.

## MATERIALS and METHODS

### Animals

The FAK^flox/flox^ strain was a generous gift from Drs. Jun-lin Guan (University of Cincinnati) and Xu Peng (Texas A&M University College of Medicine)^30^. FAK conditional KO (cKO) strain was developed by breeding *FAK^f^*^lox/flox^ strain with *K14-Cre* transgenic mice (generous gift from Dr. Elaine Fuchs, Rockefeller University)^31^, which efficiently excised floxed exons by embryonic day 15.5. Neonatal mice genotypic for K14-Cre and FAKflox/flox (cKO) were selected. All the mice were housed under pathogen-free conditions in ARC (animal resource center) of University of Chicago under a 12 hr light-dark cycle. All the subjects were not involved in any previous procedures. Mice were euthanized using controlled gradual-fill carbon dioxide (100% CO₂) from a compressed-gas cylinder, delivered through a pressure regulator and flow meter into a 7 L chamber at a displacement rate of 30–50% of the chamber volume per minute. Cervical dislocation was performed after loss of consciousness by trained personnel. Death was confirmed before tissue collection or disposal. The experimental protocol was reviewed and approved by the Institutional Animal Care and Use Committee of University of Chicago.

### Primary Epidermal Cell Culture

Primary mouse keratinocytes were isolated using previously reported methods^29^. Epidermis of newborn mice or E18.5 was separated from dermis by an overnight treatment with dispase. Then, the primary keratinocytes were dispersed from the epidermis using trypsin. Keratinocytes were co-cultured with mitomycin C-treated 3T3 fibroblast feeder cells until the third passage. Cells were maintained in E-media supplemented with 15% FBS. The final concentration of Ca^2+^ is 0.05 mM. High calcium shift was performed using E-media supplemented with 15% FBS, with Ca^2+^ at a final concentration of 1.5 mM. HEK293 cells were cultured in DMEM supplemented with 10% FBS.

### PAO and VS-6063 treatment

A 1 M stock solution of PAO was freshly prepared in 100% ethanol by warming at 37 °C for 5–10 min. Cells were treated with vehicle control, PAO (200 nM), or PAO (200 nM) together with VS-6063 (defactinib; 5 μM), diluted in culture medium. Unless otherwise indicated, PAO and VS-6063 were added simultaneously. Experiments were performed between 5 min and 2 h after treatment.

### Measurement of focal adhesion disassembly

WT or FAK KO keratinocytes were plated on fibronectin-coated dishes and transfected with plasmid encoding DsRed-Zyxin27. Transfected cells were treated with PAO (200 nM) or vehicle control immediately before imaging. Time series of images were acquired on a 3i Marianas Yokogawa-type spinning-disc confocal microscope equipped with a 63x a-plane (1.45 oil) lens and an EM charge-coupled device camera. The rate constants for focal adhesion disassembly were obtained by calculating the slope of relative fluorescence intensity decreases of individual focal adhesion on a semilogarithmic scale against time as described in our previous studies^29^.

### Focal adhesion FRAP analysis

WT or FAK KO keratinocytes were plated on fibronectin-coated 3.5-cm dishes (glass bottom) and transfected with plasmid encoding DsRed-Zyxin27. Transfected cells were treated with PAO (200 nM) before FRAP. Three prebleach events were performed followed by a 1 second bleach event. Fluorescence recovery was recorded for up to 140 seconds after photobleaching. The photokinetic experiments were analyzed using DeltaVision software to obtain t1/2 recovery (half time for fluorescence recovery) and mobile protein fractions^29^.

### Cell-cell junction FRAP analysis

WT or FAK KO keratinocytes were plated on fibronectin-coated dishes and transfected with a plasmid encoding EGFP-ZO1. Cell-cell junction formation was induced by calcium shift for 12 hours. Cells were treated with PAO (200 nM), with or without VS-6063 as indicated, and FRAP analysis was carried out as described above^35^.

### PAO-induced skin injury model

*FAK* cKO mice and wild-type littermate controls (10-12 weeks old) were treated topically with a single dose of PAO^36^. Once the mice were anesthetized, an exposure area (1.6 × 1.6 cm^2^) was marked by indelible marker on the dorsal surface of the mouse skin, and then the mice were separated into two groups (n = 3 per group). The control group was treated with vehicle (ethanol, 30 μL), whereas the experimental group was treated topically with 30 μL diluted PAO (100 μg/mouse) on the marked dorsal skin. After exposure, animals were euthanized at predefined time points for sample collection and analysis. To test whether pharmacological inhibition of FAK can affect skin vesication, VS-6063 (defactinib), an orally bioavailable FAK inhibitor, was administered to wild-type mice 30 minutes after PAO exposure (50 mg/kg).

We then analyzed skin edema and erythema. H&E and IHC analysis were used to quantify epidermal thickness, formation of microvesications, cell proliferation, apoptosis, and inflammatory cell infiltration. All histological and IHC data were analyzed and scored in blinded studies through the UChicago Pathology Core.

### Protein biochemical analysis

Western blot was conducted as previously described^37^. Cell lysates were prepared with RIPA buffer containing protease inhibitors. After the concentration of total protein is assessed, equal amounts of the cell lysates were resolved in sodium dodecyl sulfate–polyacrylamide gel and electroblotted onto a nitrocellulose membrane. The membrane was incubated with Odyssey blocking buffer (Li-COR biosciences, Lincoln, NE) for 1 h at room temperature, followed by an overnight incubation with desired primary antibody at 4°C. Immunoblots were washed three times with 1× Tween 20/phosphate-buffered saline (PBST) and incubated with secondary antibody (1:10,000 dilution) at room temperature for 1 h. Blot was washed with 1× PBST for another three times. LI-COR Odyssey scanner was used to visualize the blotting signals, and LI-COR Biosciences Software was used to conduct the quantification.

### Histology and immunofluorescence

Skin or wound samples were embedded in OCT, sectioned, and fixed in 4% paraformaldehyde. Antibodies were diluted according to manufacturer’s instruction unless indicated. Images were collected on a Zeiss AxioObserver7 and analyzed using Zeiss Zen software and ImageJ. Live cell imaging and cellular immunofluorescence images were collected using a SoRa Subdiffraction Marianas Spinning Disc Confocal. Colocalization analysis was performed using the FIJI plugin JaCoP.

## QUANTIFICATION AND STATISTICAL ANALYSIS

Statistical analysis was performed using Excel or GraphPad Prism software. Box plots are used to describe the entire population without assumptions on the statistical distribution. Student’s *t*-test was used to assess the statistical significance (*P*-value) of the difference for most experiments. *p* < 0.05 was considered statistically significant. *p* < 0.05 is indicated as one asterisk (*), and *p* < 0.01 is indicated as two asterisks (**).

## DATA AND SOFTWARE AVAILABILITY

A complete list of software for data analysis and processing can be found in the Key Resources Table.

## Acknowledgments

We thank Dr. Elaine Fuchs at the Rockefeller University, Dr. Jun-lin Guan (University of Cincinnati), and Dr. Xu Peng (Texas A&M University College of Medicine) for sharing reagents. We thank Linda Degenstein from the University of Chicago transgenic core facility for her excellent technical assistance. The animal studies were carried out in the ALAAC-accredited animal research facility at the University of Chicago.

## Data Availability Statement

For this study, we will make our data available to the scientific community, which will avoid unintentional duplication of research. All the research data will be shared openly and in a timely manner in accordance with the most recent NIH guidelines (http://grants.nih.gov/grants/policy/data_sharing/). The datasets used and/or analyzed during the current study are available from the corresponding author upon reasonable request.

## Author Contributions

XW conceived and supervised the study. HL, SS, and BT performed the experiments and analyzed the data. S-YC, MA, HL, and RA contributed to experimental design, technical expertise, and data interpretation. HL and XW wrote the manuscript. All authors reviewed and edited the manuscript.

## REFERENCES

1 Ganesan, K., Raza, S. K. & Vijayaraghavan, R. Chemical warfare agents. J Pharm Bioallied Sci 2, 166–178, doi:10.4103/0975-7406.68498 (2010).

2 Riley, B. The toxicology and treatment of injuries from chemical warfare agents. Current Anaesthesia & Critical Care 14, 149–154 (2003).

3 Nair, A. et al. Toxic blister agents: Chemistry, mode of their action and effective treatment strategies. Chem Biol Interact 350, 109654, doi:10.1016/j.cbi.2021.109654 (2021).

4 Romero, A. et al. Toxicology of Blister Agents: Is Melatonin a Potential Therapeutic Option? Diseases 9, doi:10.3390/diseases9020027 (2021).

5 Le, H. Q. & Knudsen, S. J. Exposure to a First World War blistering agent. Emerg Med J 23, 296–299, doi:10.1136/emj.2005.032540 (2006).

6 Pitschmann, V. Overall view of chemical and biochemical weapons. Toxins (Basel) 6, 1761–1784, doi:10.3390/toxins6061761 (2014).

7 Jain, A. K. et al. Histopathological and immunohistochemical evaluation of nitrogen mustard-induced cutaneous effects in SKH-1 hairless and C57BL/6 mice. Exp Toxicol Pathol 66, 129–138, doi:10.1016/j.etp.2013.11.005 (2014).

8 Shakarjian, M. P. et al. Mechanisms mediating the vesicant actions of sulfur mustard after cutaneous exposure. Toxicol Sci 114, 5–19, doi:10.1093/toxsci/kfp253 (2010).

9 Srivastava, R. K. et al. Defining cutaneous molecular pathobiology of arsenicals using phenylarsine oxide as a prototype. Sci Rep 6, 34865, doi:10.1038/srep34865 (2016).

10 Muzaffar, S., Khan, J., Srivastava, R., Gorbatyuk, M. S. & Athar, M. Mechanistic understanding of the toxic effects of arsenic and warfare arsenicals on human health and environment. Cell Biol Toxicol 39, 85–110, doi:10.1007/s10565-022-09710-8 (2023).

11 Blanpain, C. & Fuchs, E. Epidermal stem cells of the skin. Annu Rev Cell Dev Biol 22, 339–373, doi:10.1146/annurev.cellbio.22.010305.104357 (2006).

12 Proksch, E., Brandner, J. M. & Jensen, J. M. The skin: an indispensable barrier. Exp Dermatol 17, 1063–1072, doi:10.1111/j.1600-0625.2008.00786.x (2008).

13 Margadant, C., Charafeddine, R. A. & Sonnenberg, A. Unique and redundant functions of integrins in the epidermis. FASEB J 24, 4133–4152, doi:10.1096/fj.09-151449 (2010).

14 Sumigray, K. D. & Lechler, T. Cell adhesion in epidermal development and barrier formation. Curr Top Dev Biol 112, 383–414, doi:10.1016/bs.ctdb.2014.11.027 (2015).

15 Doyle, A. D., Nazari, S. S. & Yamada, K. M. Cell-extracellular matrix dynamics. Phys Biol 19, doi:10.1088/1478-3975/ac4390 (2022).

16 Watt, F. M. Role of integrins in regulating epidermal adhesion, growth and differentiation. EMBO J 21, 3919–3926, doi:10.1093/emboj/cdf399 (2002).

17 Nagano, M., Hoshino, D., Koshikawa, N., Akizawa, T. & Seiki, M. Turnover of focal adhesions and cancer cell migration. Int J Cell Biol 2012, 310616, doi:10.1155/2012/310616 (2012).

18 Srivastava, R. K. et al. Common molecular profile of multiple structurally distinct warfare arsenicals in causing cutaneous chemical vesicant injury. Sci Rep 15, 6505, doi:10.1038/s41598-024-83513-1 (2025).

19 Huveneers, S. & Danen, E. H. Adhesion signaling - crosstalk between integrins, Src and Rho. J Cell Sci 122, 1059–1069, doi:10.1242/jcs.039446 (2009).

20 Mitra, S. K., Hanson, D. A. & Schlaepfer, D. D. Focal adhesion kinase: in command and control of cell motility. Nat Rev Mol Cell Biol 6, 56–68, doi:10.1038/nrm1549 (2005).

21 Mitra, S. K. & Schlaepfer, D. D. Integrin-regulated FAK-Src signaling in normal and cancer cells. Curr Opin Cell Biol 18, 516–523, doi:10.1016/j.ceb.2006.08.011 (2006).

22 Parsons, S. J. & Parsons, J. T. Src family kinases, key regulators of signal transduction. Oncogene 23, 7906–7909, doi:10.1038/sj.onc.1208160 (2004).

23 Wadhawan, A., Smith, C., Nicholson, R. I., Barrett-Lee, P. & Hiscox, S. Src-mediated regulation of homotypic cell adhesion: implications for cancer progression and opportunities for therapeutic intervention. Cancer Treat Rev 37, 234–241, doi:10.1016/j.ctrv.2010.08.003 (2011).

24 Oetken, C. et al. Phenylarsine oxide augments tyrosine phosphorylation in hematopoietic cells. Eur J Haematol 49, 208–214, doi:10.1111/j.1600-0609.1992.tb00048.x (1992).

25 Shekels, L. L., Smith, A. J., Van Etten, R. L. & Bernlohr, D. A. Identification of the adipocyte acid phosphatase as a PAO-sensitive tyrosyl phosphatase. Protein Sci 1, 710–721, doi:10.1002/pro.5560010603 (1992).

26 Zhang, Z. Y., Davis, J. P. & Van Etten, R. L. Covalent modification and active site-directed inactivation of a low molecular weight phosphotyrosyl protein phosphatase. Biochemistry 31, 1701–1711, doi:10.1021/bi00121a018 (1992).

27 Pelham, R. J., Jr. & Wang, Y. Cell locomotion and focal adhesions are regulated by substrate flexibility. Proc Natl Acad Sci U S A 94, 13661–13665, doi:10.1073/pnas.94.25.13661 (1997).

28 Retta, S. F. et al. Focal adhesion and stress fiber formation is regulated by tyrosine phosphatase activity. Exp Cell Res 229, 307–317, doi:10.1006/excr.1996.0376 (1996).

29 Wu, X., Kodama, A. & Fuchs, E. ACF7 regulates cytoskeletal-focal adhesion dynamics and migration and has ATPase activity. Cell 135, 137–148, doi:S0092-8674(08)01011-8 [pii]10.1016/j.cell.2008.07.045 (2008).

30 Shen, T. L. et al. Conditional knockout of focal adhesion kinase in endothelial cells reveals its role in angiogenesis and vascular development in late embryogenesis. J Cell Biol 169, 941–952, doi:jcb.200411155 [pii]10.1083/jcb.200411155 (2005).

31 Vasioukhin, V., Degenstein, L., Wise, B. & Fuchs, E. The magical touch: genome targeting in epidermal stem cells induced by tamoxifen application to mouse skin. Proc Natl Acad Sci U S A 96, 8551–8556 (1999).

32 Banerjee, S. et al. Defactinib with avutometinib in patients with solid tumors: the phase 1 FRAME trial. Nat Med 31, 3074–3080, doi:10.1038/s41591-025-03763-y (2025).

33 Lin, H. M. et al. Effect of FAK inhibitor VS-6063 (defactinib) on docetaxel efficacy in prostate cancer. Prostate 78, 308–317, doi:10.1002/pros.23476 (2018).

34 Hoffman, R. D. & Lane, M. D. Iodophenylarsine oxide and arsenical affinity chromatography: new probes for dithiol proteins. Application to tubulins and to components of the insulin receptor-glucose transporter signal transduction pathway. J Biol Chem 267, 14005–14011 (1992).

35 Ma, Y. et al. ACF7 regulates inflammatory colitis and intestinal wound response by orchestrating tight junction dynamics. Nat Commun 8, 15375, doi:10.1038/ncomms15375 (2017).

36 Srivastava, R. K. et al. Unfolded protein response (UPR) signaling regulates arsenic trioxide-mediated macrophage innate immune function disruption. Toxicol Appl Pharmacol 272, 879–887, doi:10.1016/j.taap.2013.08.004 (2013).

37 Wu, X., Gan, B., Yoo, Y. & Guan, J. L. FAK-mediated src phosphorylation of endophilin A2 inhibits endocytosis of MT1-MMP and promotes ECM degradation. Dev Cell 9, 185–196, doi:S1534-5807(05)00249-2 [pii]10.1016/j.devcel.2005.06.006 (2005).

