## supplementary information for "FAK-Dependent Focal Adhesion Remodeling Drives Arsenical Vesicant–Induced Skin Injury"

**Supplementary Figure 1. Basal cell-cell junction dynamics in WT and FAK-null keratinocytes.** Representative FRAP sequences of EGFP-ZO1 at cell-cell junctions in untreated (A) WT and (B) FAK-null keratinocytes. Enlarged regions are indicated by yellow boxes. Numbers indicate seconds after photobleaching.

**Supplementary Figure 2. Quantification of PAO-induced inflammatory and epidermal responses in WT and FAK cKO skin.** Quantitative analysis of (A) COX-2-positive area, (B) p63-positive nuclei, (C) Ki67-positive nuclei, and (D) phosphorylated NF-κB p65 (p-p65)-positive nuclei in skin from WT and epidermal-specific FAK conditional knockout (cKO) mice following vehicle control or PAO treatment. Error bars represent mean ± SD; n= 50 (fields). *: p<0.05, **: p<0.01 (Student’s t-test).

**Supplementary Figure 3. Epidermal FAK deficiency attenuates PAO-induced cell death.** (A) Representative TUNEL staining of skin from WT and epidermal-specific FAK conditional knockout (cKO) mice following vehicle control (Ctrl) or PAO treatment. Scale bars, 50 μm. (B) Quantification of TUNEL-positive nuclei. Error bars represent mean ± SD; n= 50 (fields). *: p<0.05, **: p<0.01 (Student’s t-test).

**Supplementary Figure 4.** Pharmacological inhibition of FAK preserves cell-cell junctions following PAO treatment. Representative immunofluorescence staining of E-cadherin (green) and DAPI (blue) in PAO-treated WT keratinocytes in the absence or presence of VS-6063. Scale bar, 50 μm.

**Supplementary Figure 5.** AlphaFold predicted binding mode between human FAK1 and PTPN12 catalytic domain (residue 1-315) (**A**). The pTyr397 residue of FAK is positioned right into the active site of PTPN12, with the phosphate group positioned right next to the catalytic Cys231 residue (**B**). PAO is known to target adjacent cysteine residues in proteins. Right next to PTPN12 Cys231 is Cys235. These two cysteine residues are the likely target site of PAO, which would inhibit PTPN12 catalytic activity, leading to increased FAK1 pTyr397.

**Supplementary Figure 6. Original uncropped western blot images.** Images correspond to Figure 1C and show phosphorylated FAK (Tyr397), total FAK, and actin in control (Ctrl) and PAO-treated samples. Actin served as the loading control. Molecular weight markers are indicated in kilodaltons (kDa).

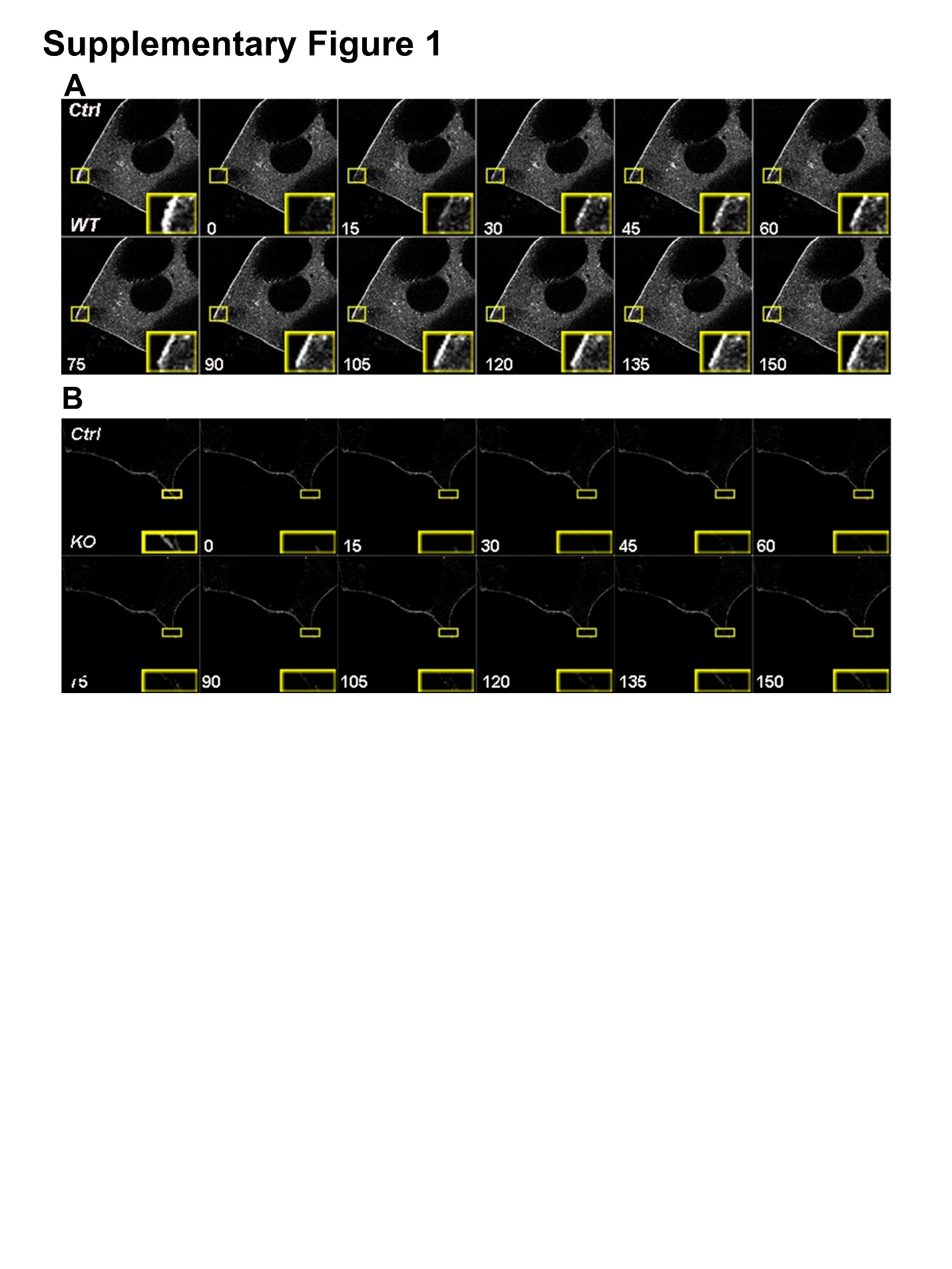

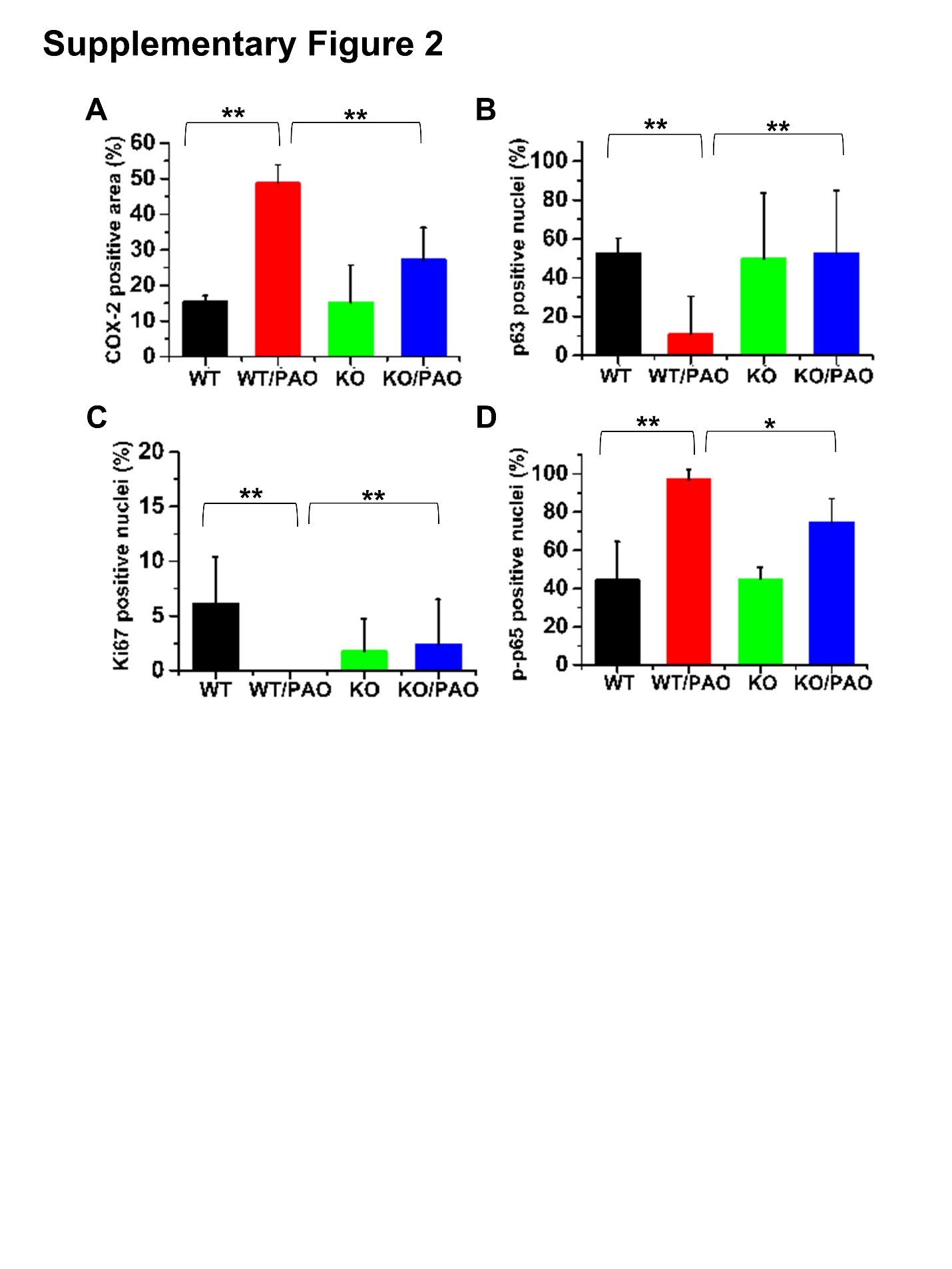

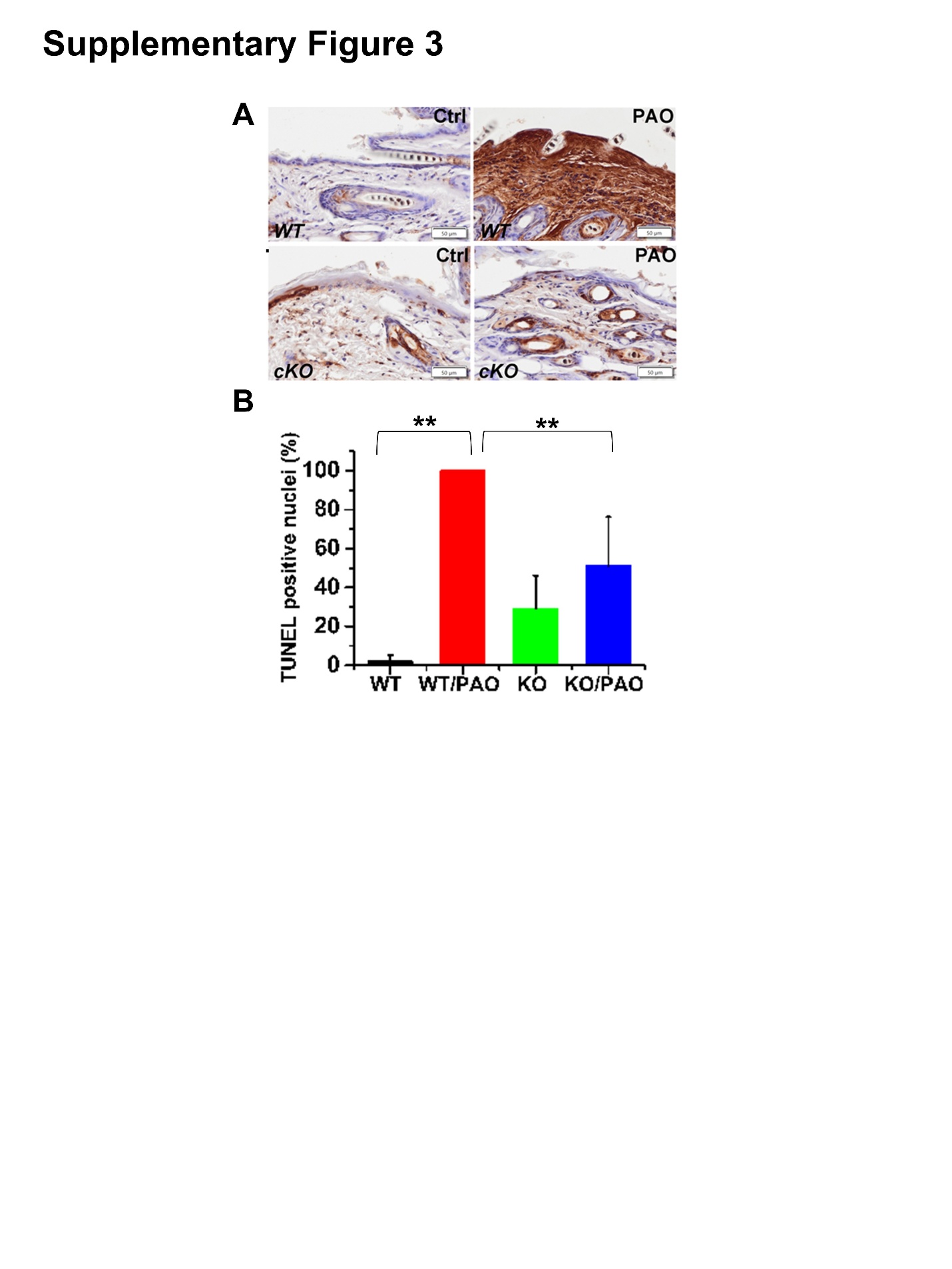

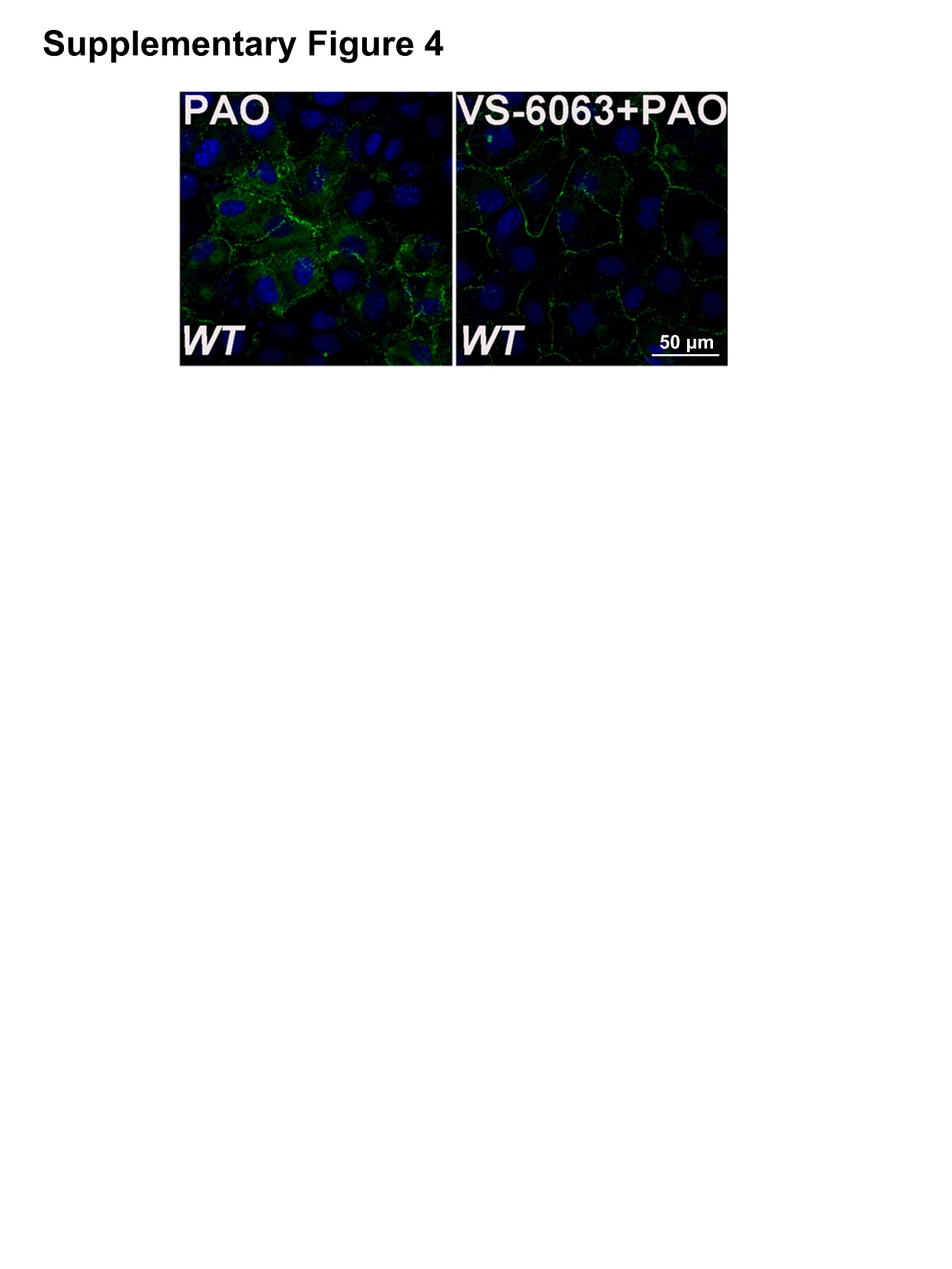

**Supplementary Figure 5**

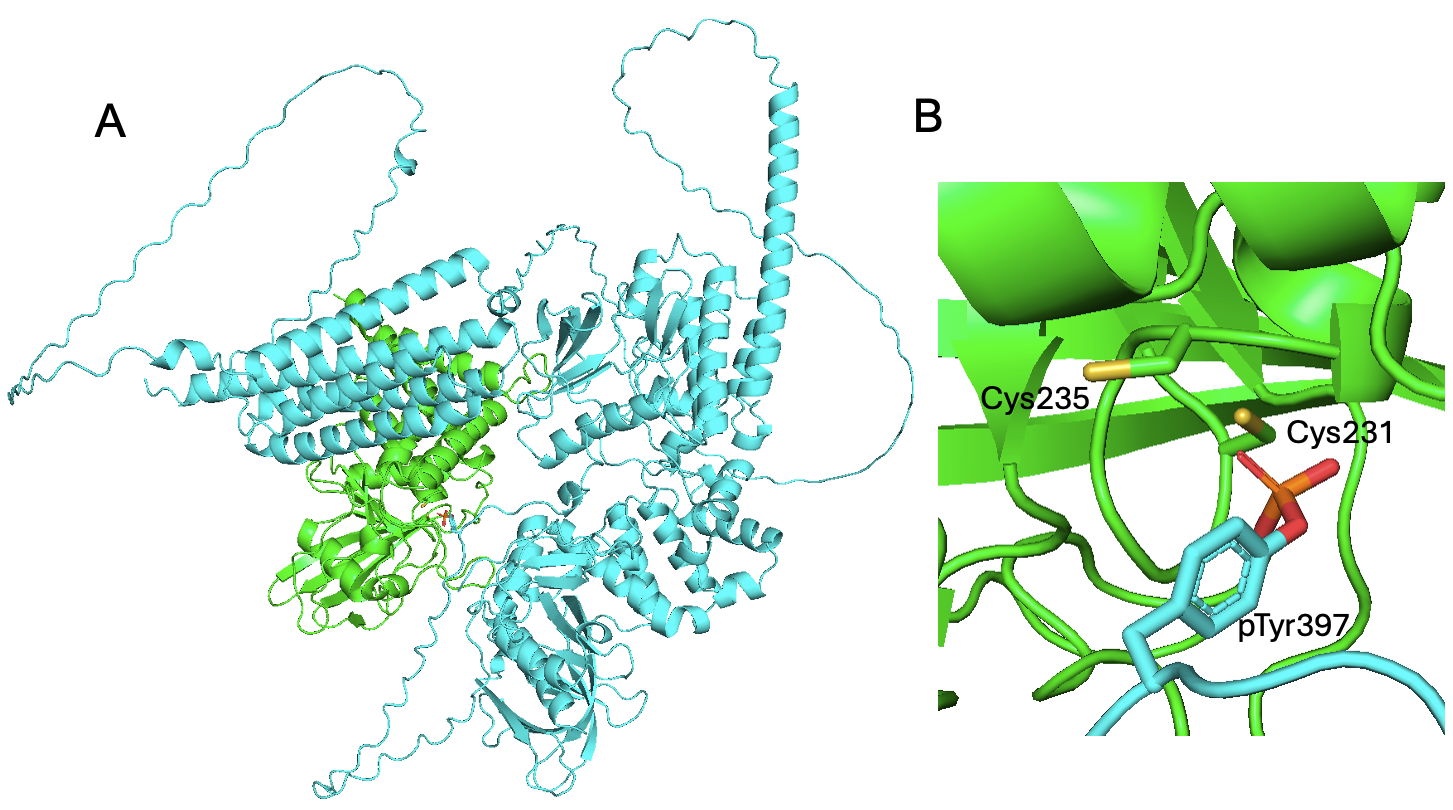

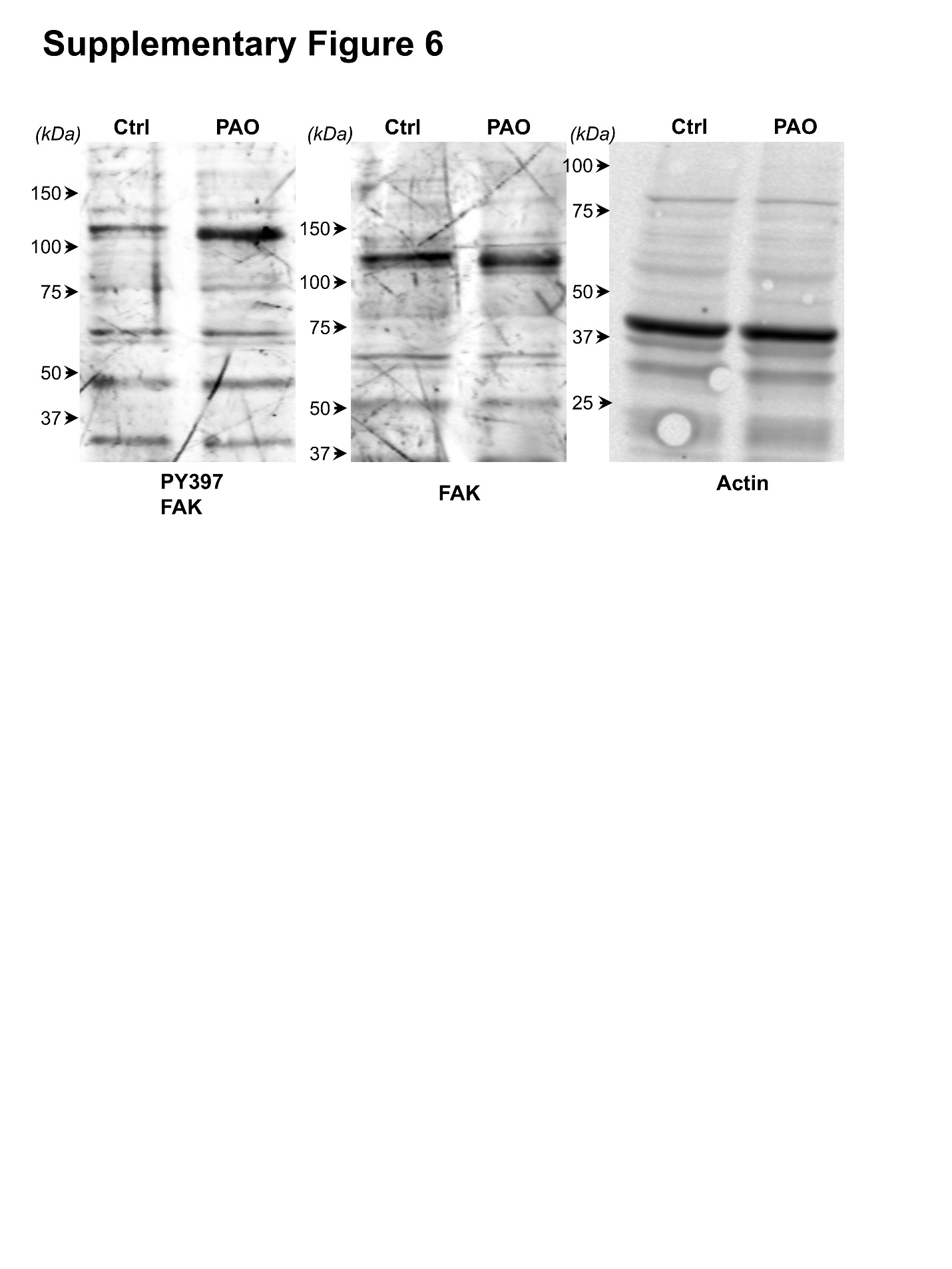
